# Computational pathology infers clinically relevant protein levels and drug response in breast cancer by weakly supervised contrastive learning

**DOI:** 10.1101/2023.04.13.536819

**Authors:** Hui Liu, Xiaodong Xie, Bin Wang

**Affiliations:** College of Computer and Information Engineering, Nanjing Tech University, Nanjing, 211816, Jiangsu, China; Department of Cardiothoracic Surgery, the Third Affiliated Hospital of Soochow University, Changzhou, 213110, Jiangsu, China

**Keywords:** Weakly-supervised learning, Computational pathology, Contrastive learning, Drug response, Protein level

## Abstract

Visual inspection of histopathology slides via optical microscope is the routine medical examination for clinical diagnosis of tumors. Recent studies have demonstrated that computational pathology not only automate the tumor diagnosis, but also showed great potential to uncover tumor-related genetic alterations and transcriptomic patterns. In this paper, we propose wsi2rppa, a weakly supervised contrastive learning framework to infer the protein levels of tumor biomarkers from whole slide images (WSIs) in breast cancer. We firstly conducted contrastive learning-based pre-training on tessellated tiles to extract histopathological features, which are then aggregated by attention pooling and adapted to downstream tasks. Our extensive experiments showed that our method achieved state-of-the-art performance in tumor diagnostic task, and also performed well in predicting clinically relevant protein levels and drug response. To show the model interpretability, we spatially visualized the WSIs colored the tiles by their attention scores, and found that the regions with high scores were highly consistent with the tumor and necrotic regions annotated by a 10-year experienced pathologist. Moreover, spatial transcriptomic data further verified that the heatmap generated by attention scores agree greatly with the spatial expression landscape of two typical tumor biomarker genes. In particular, our method achieved 0.79 AUC value in predicting the response of breast cancer patients to the drug trastuzumab treatment. These findings showed the remarkable potential of deep learning-based morphological feature is very indicative of clinically relevant protein levels, drug response and clinical outcomes.

## 1 Introduction

Digital pathology makes use of automated microscopy or optical magnification systems to scan and digitize traditional glass pathology slides. Computer algorithms are then employed to perform high-precision multi-field seamless stitching and processing of the images to produce whole-slide images (WSIs). In recent years, computational pathology has advanced rapidly to facilitate the automatic diagnosis of tumors. This not only alleviates the workload of pathologists, but also helps to eliminate the subjective bias from observers.

Deep learning has made significant strides in many fields, and has also spurred advances in computational pathology. Such methods have demonstrated impressive progress in a variety of challenging clinical tasks [1–4]. For tumor diagnosis and semantic segmentation, Wang et al. [5] developed an enhanced HD-Staging algorithm based on mask-RCNN that can perform nuclear segmentation and cell classification on lung adenocarcinoma images. This algorithm can also be applied to head and neck cancer, breast cancer, and lung squamous cell cancer. Greenwald et al.[6] developed TissueNet to segment cells and identify the boundaries of individual cells in whole-slide images. Eklund et al.[7] created an integrated model based on Inception V3 to assist pathologists in finding, identifying, and grading prostate tumors. Wang et al.[8] introduced a novel deep learning model to improve breast cancer histological grading. Additionally, several studies have presented computational pathology methods for inferring genetic alterations and gene expression profiles. For example, Kather and colleagues [9] introduced a deep learning approach for analyzing genetic alterations across various types of cancer. They successfully predicted the presence of CDC27 mutations in colorectal cancer using pathological images. Wang et al.[10] proposed an approach called expression-morphology (EMO) to predict mRNA expression in 17,695 genes from whole-slide images and validated the spatial variability of intratumor expression through spatial transcriptomics profiling. Several recent studies have demonstrated the potential of using weakly supervised algorithms in computational pathology. For instance, our team has developed Histcode [11], which integrated contrastive learning-based pretraining and weakly supervised learning to predict differential gene expression of cancer driver genes. Mahmood et al.[12] developed the CLAM model, which used an attention mechanism for subtype classification of renal cell carcinoma and non-small cell lung cancer. Schumach et al.[13] developed a model called HE2RNA to predict RNA-seq from pathological images and identified tumors with microsatellite instability in clinical diagnoses. Shamai et al.[14] built convolutional neural network (CNN)-based model to predict PD-L1 expression from H&E images. Eliceiri et al. [15] developed the DS-MIL algorithm to classify WSIs as tumor or normal. Mahmood et al. [16] developed SISH, an interpretable search pipeline that achieves high speed in searching histology images after training with only slide-level labels.

In recent times, the field of proteomics has demonstrated exceptional promise in the realm of precision medicine, achieved significant advances in the treatment of various forms of cancer [17–19]. For example, Gao et al. [20] utilized proteomics to identify two prognostic biomarkers, PYCR2 and ADH1A, for hepatocellular carcinoma. The reverse-phase protein array (RPPA) [21], a high-throughput and highly sensitive protein microarray that employs antibodies to detect and quantify proteins and their modifications in biological samples, has shown tremendous potential in the discovery of cancer biomarkers, analysis of functional phenotypes, and elucidation of drug mechanisms. It has been assumed that protein profile changes in tumor cells cause functional changes, which can influence tumor cell morphology. So, routine histopathology tissue slides, which are ubiquitously available, can reflect such morphological changes, thereby the clinically relevant proteins could be inferred directly from digitized whole-slide images.

In this paper, we introduced a weakly supervised contrastive learning methods to establish the quantitative associations between clinically relevant proteins and histopathological features in breast cancer. Specifically, whole-slide images were split into tiles and then MoCo v2 was employed to extract features at the tile-level. Attentive pooling was used to aggregate tile-level features to generate slide latent representations, which were then applied to various downstream tasks, including tumor diagnosis, quantification of protein levels, prognostic risk assessment, and prediction of response to the drug trastuzumab. Our extensive experiments showed that the proposed method achieved state-of-the-art perfomance in tumor diagnosis task, and achieved high performance in quantifying the protein levels of tumor biomarkers. To show the model interpretability, we spatially visualized the WSIs colored by attention scores of tiles, and found that the regions with high scores are highly consistent with the tumor and necrotic regions annotated by an experienced pathologist. Moreover, spatial transcriptomic data further verified that the heatmaps generated by attention scores agree greatly with the spatial expression map of tumor biomarker genes. Our method achieved 0.79 AUC value in predicting the response of breast cancer patients to the drug trastuzumab. These findings showed our method could effectively elucidate and quantify genotype–phenotype links in breast cancer.

## 2 Results

### 2.1 Weakly-supervised contrastive learning framework

Our learning framework consisted of four steps, as illustrated in Figure 1. The first step is the preprocessing of whole slide images (WSIs). We eliminated the regions without sufficient pathological tissue and background, and then split each slide into 256*256px tiles, yielding a total of 17,020,990 tiles with an average of 8,706 tiles per slide.

**Fig. 1:**
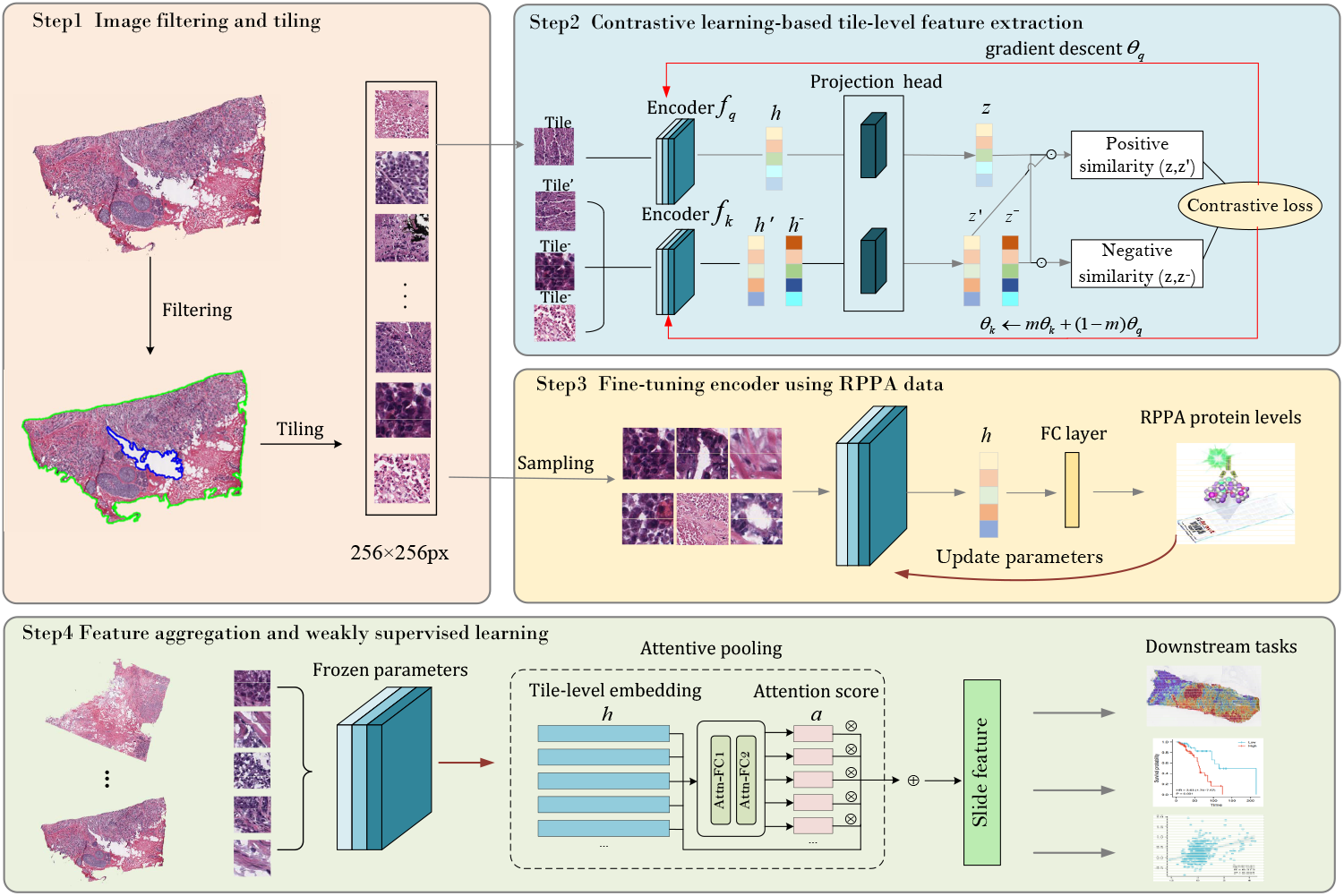
Illustrative flowchart of the proposed wsi2rppa pipeline. The first step preprocessed the WSIs by eliminating regions without sufficient tissue and splitting the slide into tiles. The second step leveraged MoCo v2 contrastive learning to extract tile-level features. The third step tuned the pre-trained encoder using protein levels measured by RPPA assays. Finally, the tile-level features were aggregated using attentive pooling to obtain slide-level features for downstream tasks.

The second stage is the contrastive learning-based feature extraction for unlabeled tiles. Contrastive learning employs self-supervised pretext tasks to learn image embedding and has demonstrated exceptional performance on quite a few tasks. We evaluated two typical contrastive learning algorithms, MoCo v2 and SimCLR, and found that they learned more informative features in comparison to the ResNet50 baseline. In the third stage, we utilized protein expression levels measured by Reverse Phase Protein Array (RPPA) assays to fine-tune the pre-trained encoder. Due to the too large number of tiles, we sampled 20% tiles (m=398,426) from 200 randomly selected slides to run the fine-tuning task. Finally, we employed attentive pooling to aggregate tile-level features into slide feature ready for downstream tasks, including tumor diagnosis, prediction of biomarker gene expression levels and drug treatment outcome, and establishment of prognostic score.

### 2.2 Weakly-supervised learning generate accurate tumor diagnosis

We first tested our method for tumor diagnostic task on the TCGA-BRCA cohort. The pathology images (n=1,979, see Section 3.1) of breast cancer were randomly split into training, validation and independent test set by 60%, 20% and 20%. It is worth noting that we used only the slide-level labels to finetune the pretrained model. Our method achieved 0.995 accuracy and 0.996 AUC values. As shown in Figure 2 (a-b), we presented the ROC curve and confusion matrix on the independent test set. This showed that our weakly supervised learning model can accurately discriminate whether WSI contains tumor cells or not. During the fine-tuned stage for tumor diagnosis, the tile-level embeddings were aggregated to build slide representation (computational histopathological features). As the network was trained to classify tumor and normal tissues, the computational histopathological features showed two clusters corresponding to each tissue class by PCA dimension reduction (Figure 2 (c)).

**Fig. 2:**
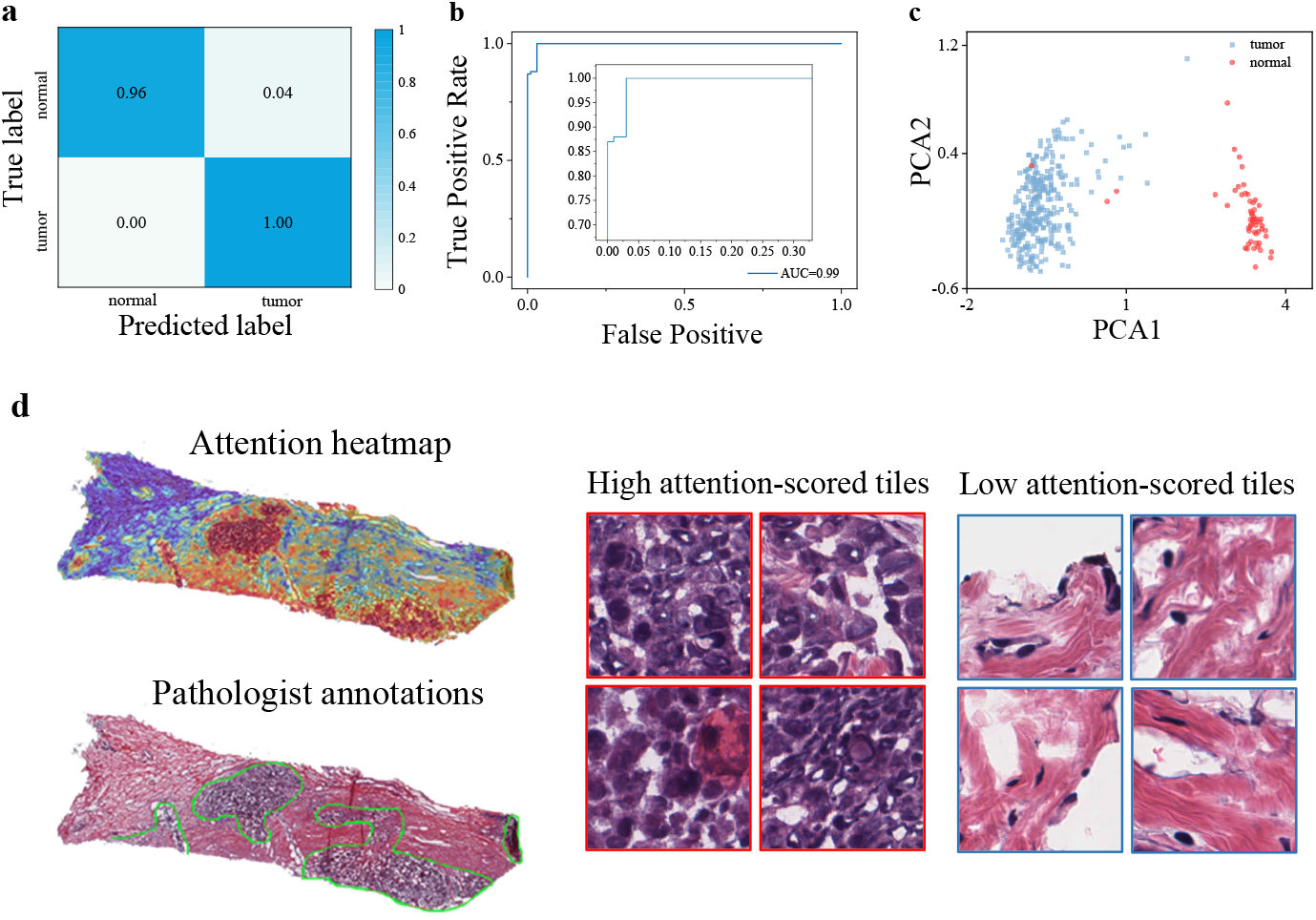
**(a-b)** The confusion matrix and ROC curve for tumor diagnosis task on the test set of TCGA-BRCA cohort. **(c)** Visualization of learned slide-level feature by PCA dimension reduction. **(d)**Comparison between the heatmap generated by tile-level attention scores and the tumor area annotated by a pathologist. The heatmap (upper left) is generated by the spatial deconvolution of the tiles to original slide (lower left), coloring each tile according to its attention score. The area circled by the green line is the tumor necrosis area depicted by an experienced pathologist (lower left). The right part shows a few representative tiles with the highest and lowest attention scores.

The slide-level classification of tumor/normal labels was made based on the attentive pooling of tile-level features, thereby the derived attention scores are highly indicative of localization of tumor regions. To validate the spatial localization of tumor area by our model, we scaled the learned tile-level attention scores to generate heatmaps by spatial deconvolution of tiles to original slide. For objective evaluation, an experienced pathologist was asked to annotate the tumor regions of interest (ROIs), and we visually checked whether the high-scored area was coincident the annotated regions. As shown in Figure 2 (d), the attention heatmaps showed remarkable agreement between the pathologist-annotated ROIs and our computational predictions of tumor regions. Visual inspection of tiles would reflect more human-readable pathological feature. So, we showed a few tiles with high or low attention scores, which are verified by pathologist that the high-scored tiles are mostly normal tissue, while high-scored tiles are tumor tissue.

### 2.3 Histopathological feature effectively predicts protein levels of tumor biomarkers

The protein expression profiles of clinically relevant biomarkers produced by reverse-phase protein arrays (RPPAs) allow us to test whether the histopathological feature is predictive of protein levels of tumor biomarkers. As these proteins are enriched in diagnostic biomarkers and therapeutic targets, establishment of the quantitative associations between computational pathology and clinically relevant proteins would greatly facilitate clinically translational applications. On the TCGA-BRCA cohort, the protein expression profile contains 223 biomarker proteins of 860 breast cancer patients. We transferred the computational pathological features obtained by contrastive learning to quantitatively predict the levels of biomarker proteins.

We need to make it clear that we used multi-task learning to simultaneously predict the expression level of 223 biomarker proteins. This is quite different from prior methods that trained a predictive model per gene or genetic mutant [9, 10, 22]. For evaluation of this regression task, we used Pearson correlation coefficient (*r*) between the RPPA-measured values and predicted values as the evaluation metric. Similar to previous studies [13], we considered a prediction to be significantly different from the random baseline value if the *p*-value associated with its coefficient was below 0.05, after applying Benjamini–Hochberg (BH) correction to account for multiple-hypothesis testing. On the TCGA-BRCA cohort, almost all proteins yield significant prediction results. As shown in Figure 3 (a-b), 220 out of 223 protein (99%) were predicted with a statistically significant correlation under BH correction. The average correlation coefficient reached 0.292, of which 164 proteins obtained correlation coefficient greater than 0.2, and 16 proteins greater than 0.5.

**Fig. 3:**
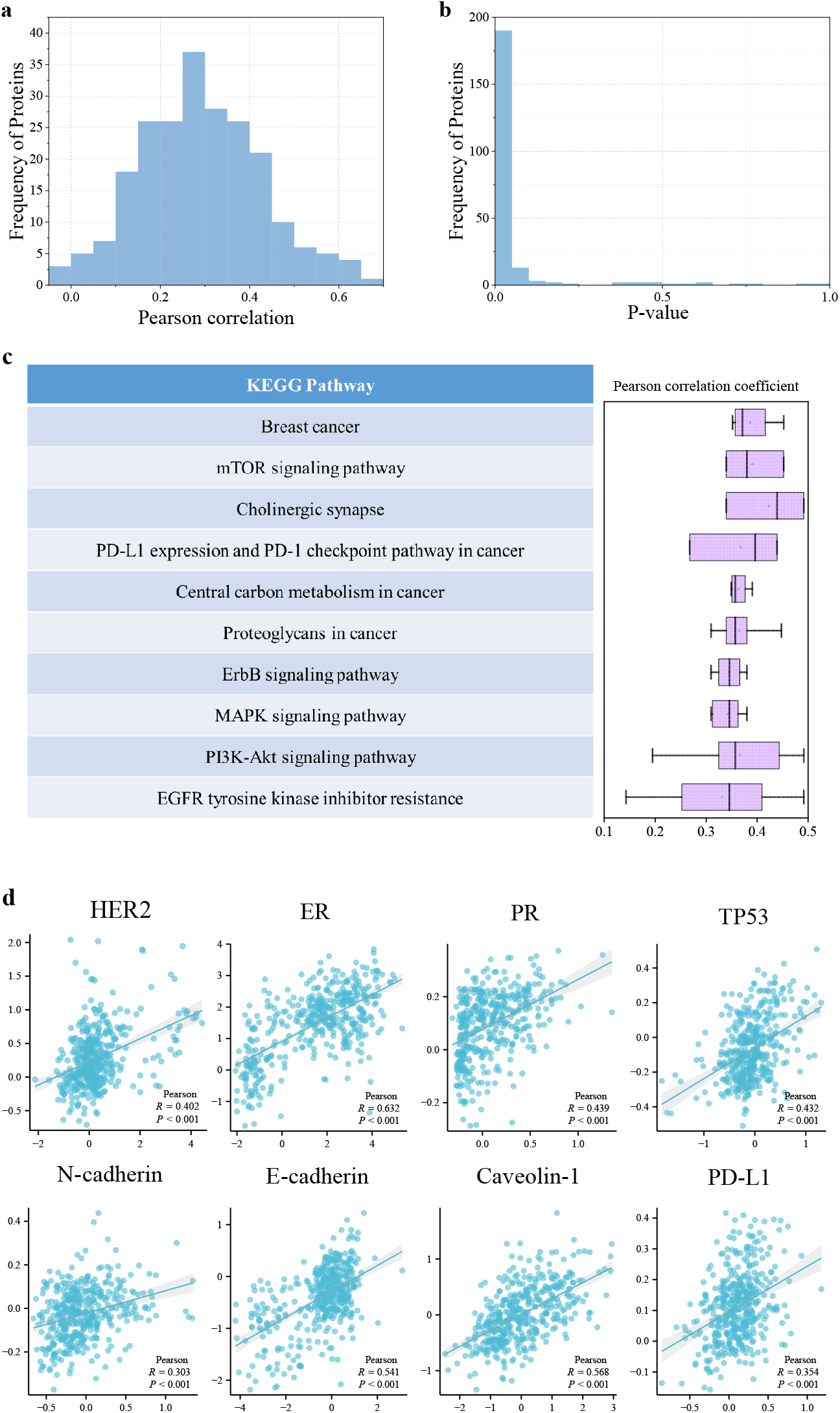
**(a-b)** showed the histogram of the Pearson correlation coefficients between predicted values and true values of 223 clinically relevant proteins, and the histogram of *p*-values. **(c)** showed the KEGG pathways the one third of well-predicted proteins enriched in (left), and the boxplots of the Pearson coefficients of proteins involved in each signal pathway (right). (d) Scatter plots of the proteins closely associated to targeted therapy and immunotherapy in breast cancer.

Among the tumor biomarkers, we paid close attention to the proteins that are clinically relevant to the breast cancer, such as estrogen receptor (ER), progesterone receptor (PR) and HER2. Expectedly, our model showed strong predictive power on these biomarkers, as shown in Figure 3 (d). For the estrogen receptor ER-*α*, positive in 70% of breast cancer and key biomarker for breast cancer diagnosis [23], our method got *r*=0.632 correlation coefficient. Progesterone receptor (*r*=0.439) is induced by ER-*α* and plays an important role in regulating ER-*α* protein, thereby it functions as an important biomarker for breast cancer treatment and prognosis [24]. Highly expressed PR in luminal A type breast cancer indicates good prognosis [25]. HER2 (*r*=0.402) overexpression is present in 20-30% of breast cancer and is associated with higher malignancy [26]. The tumor suppressor factor TP53 (*r*=0.432) has mutated in 80% of triple-negative breast cancers [27]. Caveolin-1 (*r*=0.568) high expression is a biomarker of more malignant tumor and poor prognosis [28]. PD-L1 (*r*=0.354) is an immune-related biomarker expressed on the surface of several cell types. Its high expression means potential response to immunotherapy, specifically PD-1/PD-L1 immune checkpoint inhibitors [29]. E-cadherin (*r*=0.541) and N-cadherin (*r*=0.303) are epithelial–mesenchymal transition (EMT)-related biomarkers that confer higher invasion, metastasis and drug resistance [30]. Moreover, we choose one third well-predicted proteins (n=75) to run functional enrichment analysis, and the result was shown in Figure 3 (c), from which we found that the well-predicted protein are enriched in the oncogenic, drug-resistance and tumor-driven signaling pathways.

### 2.4 Contrastive learning significantly improves predictive performance

To verify that the pre-trained encoder by contrastive learning extracted histologic features generalized to various downstream tasks, we compared two popular contrastive learning models (SimCLR and MoCo v2) to the baseline ResNet50 network. The ResNet50 was trained on ImageNet and transferred to the prediction task of protein levels. The contrastive learning used the ResNet50 as backbone network. When the contrastive learning-based pretraining finished, the ResNet50 encoder was frozen in the transfer learning for protein levels prediction task. For comprehensive comparison, we also considered fine-tuning the pre-trained ResNet50 encoder by appending a fully-connected layer to predict the protein levels. As shown in Figure 4 (a), all the model pretrained by contrastive learning outperformed the baseline ResNet50 model. Also, we found that fine-tuned models achieved better predictive power than their respective frozen ones. The experimental result demonstrated the contrastive learning effectively capture expressive and informative features that can be transferred to improve performance of downstream tasks.

**Fig. 4:**
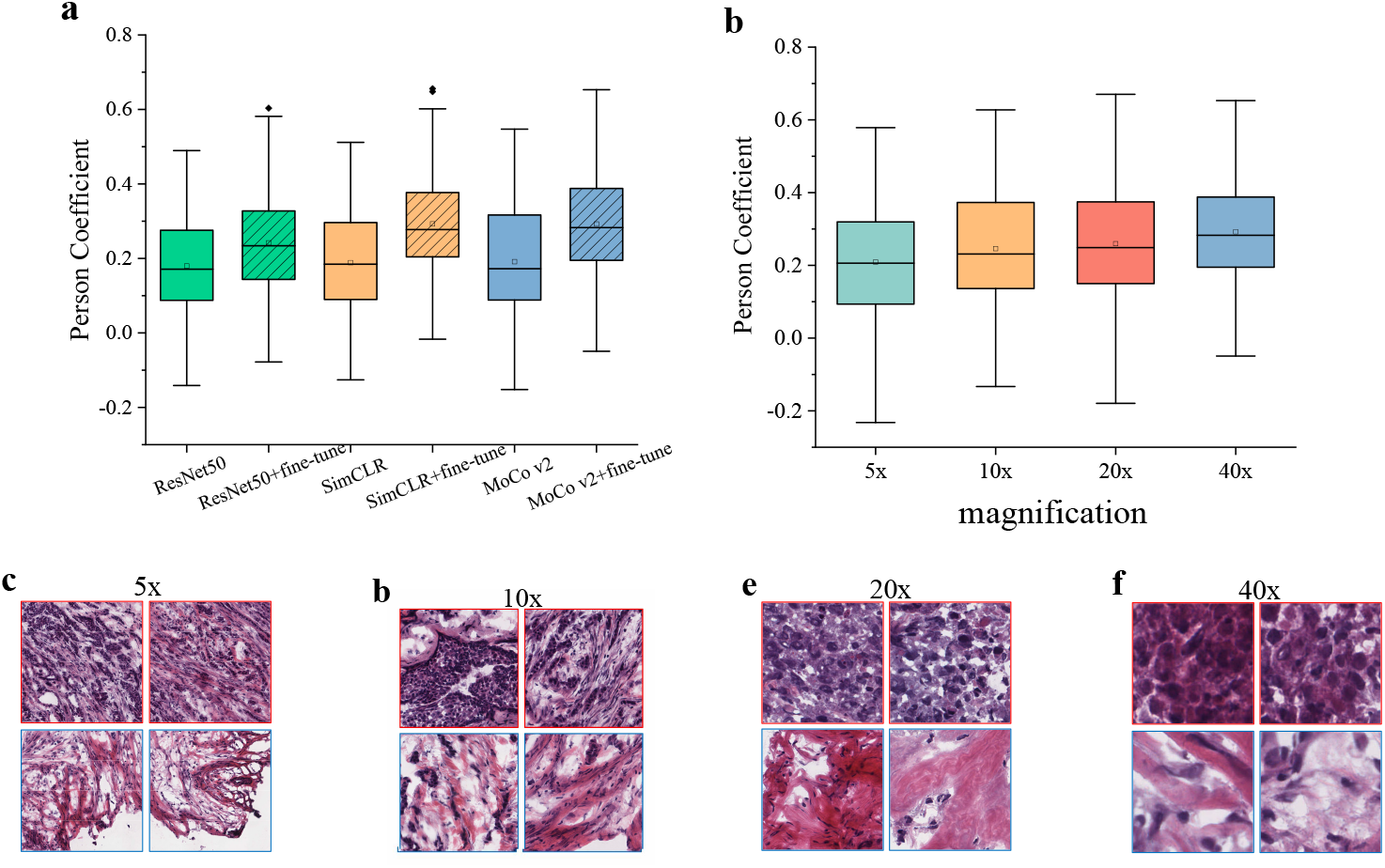
**(a)** Boxplots of the Pearson correlation coefficients between the real and predicted protein levels using the tile features extracted by ResNet50, Sim-CLR, MoCo v2, as well as their fine-tuned versions, respectively. **(b)** Boxplots of the Pearson correlation coefficients acquired using the tiles at magnification of 5x, 10x, 20x and 40x. **(c-f)** show some representative tiles at magnification of 5x, 10x, 20x, 40x.

We were also interested in the effect of different resolution of the scanned pathology images on the predictive performance. We have tested the 40x, 20x, 10x and 5x magnification, and the experimental results shows that higher magnification achieved better performance in the prediction of protein levels. Figure 4 (b) showed the boxplots of correlation coefficients at different resolutions, the 40x, 20x, 10x and 5x magnification yield 0.292, 0.259, 0.246 and 0.209, respectively. For intuitive proof, we visually checked a few exemplar tiles from different resolution slides, as shown in Figure 4 (c-f) and found that higher resolution slides convey more detailed information about the morphological structure and histopathological feature. We speculated that higher magnification enable the model to better capture the molecular phenotypic feature of pathological tissues.

### 2.5 Computational pathological feature indicates patient prognosis

After aggregation of tile-level features by attentive pooling strategy, we obtained the latent representation of a whole slide image. Following the feature aggregation, we further tested whether the computational pathological feature is indicative of the prognosis of breast cancer patients. For this purpose, a Cox proportional hazards regression model was developed to estimate the survival risk (see Section 3.10). According to the established prognostic risk score, the breast cancer patients (n=860) of TCGA-BRCA cohort were stratified to high-risk group and the low-risk group, and the survival analysis curve was shown in Figure 5 (a). The result showed that the 5-year survival time of the high-risk group was significantly lower than that of the low-risk group (HR=3.63, p-value*<*0.05), and the concordance index (C-index) of the survival model was 0.64. Moreover, we compared the histopathological difference of the high-risk and low-risk pathology images. As shown in Figure 5 (b), it can be found that the high-risk slide contains more tumor regions (high attention score regions in the heatmap) and conveys more invasive histologic feature, compared to the low-risk slide. These results showed that our method automatically capture histologic feature that was indicative of patient prognosis, which was different from other methods based on manually annotated ROIs.

**Fig. 5:**
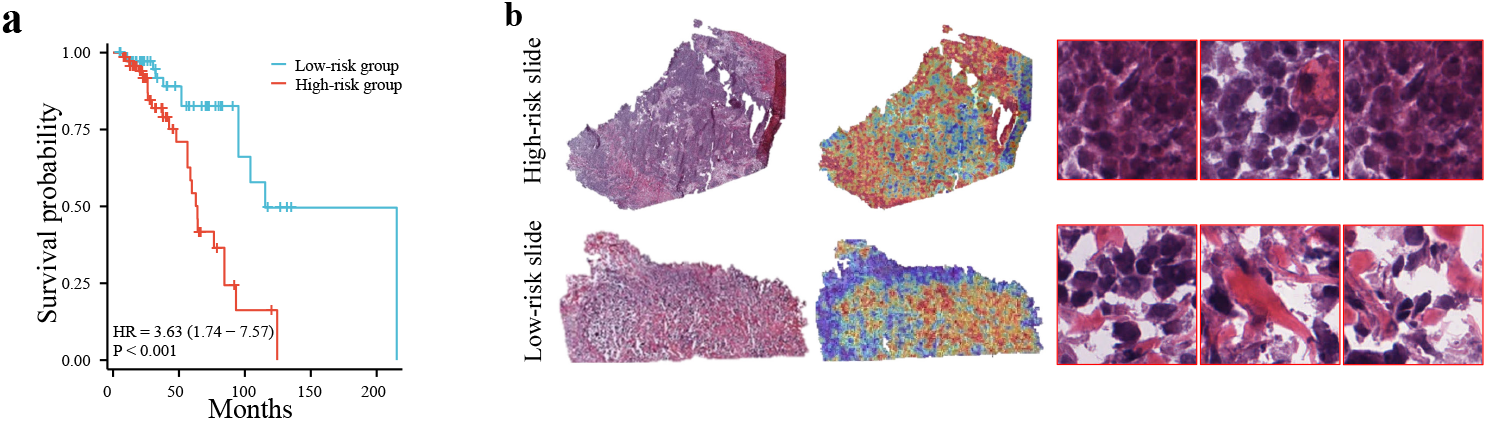
**(a)** Survival analysis for the stratified breast cancer patients of TCGA-BRCA cohorts using the risk scores based on computational pathological features. **(b)** shows two representative pathology images with high-risk score and low-risk score and their corresponding attention heatmaps, as well as some exemplar tiles.

### 2.6 Computational pathology predicts trastuzumab treatment outcome

We went further to explore the capacity of computational pathology in predicting drug response of breast cancer. The Yale trastuzumab response cohort [31] allows us to run this exploratory experiment. This cohort included the patients with a pre-treatment breast core biopsy with HER2 positive invasive breast carcinoma who then received neoadjuvant targeted therapy with trastuzumab *±*pertuzumab prior to definitive surgery. A case was designated as responder if the pathological examination of surgical resection specimens did not report residual invasive, lympho-vascular invasion or metastatic carcinoma, and otherwise non-responders. After removal of the H&E images without enough detectable foreground tissue by redefined area threshold, we obtained 75 cases (34 responders and 41 non-responders) from the Yale trastuzumab response cohort.

As the HER2 status is used as the clinical biomarker for trastuzumab response of breast cancer, we used only the RPPA-level of HER2, rather than the clinical treatment outcomes, to fine-tune the model pretrained by contrastive learning. Thereafter, the fine-tuned model was directly used to predict the trastuzumab treatment response. This would empower our method to approach clinical practice and output actionable suggestion. As shown in Figure 6 (a-c), our method achieved 0.79 AUC value for the prediction of responder and non-responders, which is much better than 0.68 obtained by the previous study [31]. Of note, our model was trained by unannotated slides, but achieved comparable performance to previous model trained using pathologist-annotated slides with invasive tumor cells area. Again, we visualized a heatmap generated by the learned attention scores, and compared the highly scored regions to the tumor regions annotated by the pathologists of Yale University. Figure 6 (d) demonstrated high consistence between them, and the tiles randomly picked out from high- and low-scored regions agree remarkably with the regions marked as normal and tumor. These results supported the feasibility of image-based biomarkers to predict trastuzumab treatment and the ability of deep learning model to recognize morphological changes related to treatment results.

**Fig. 6:**
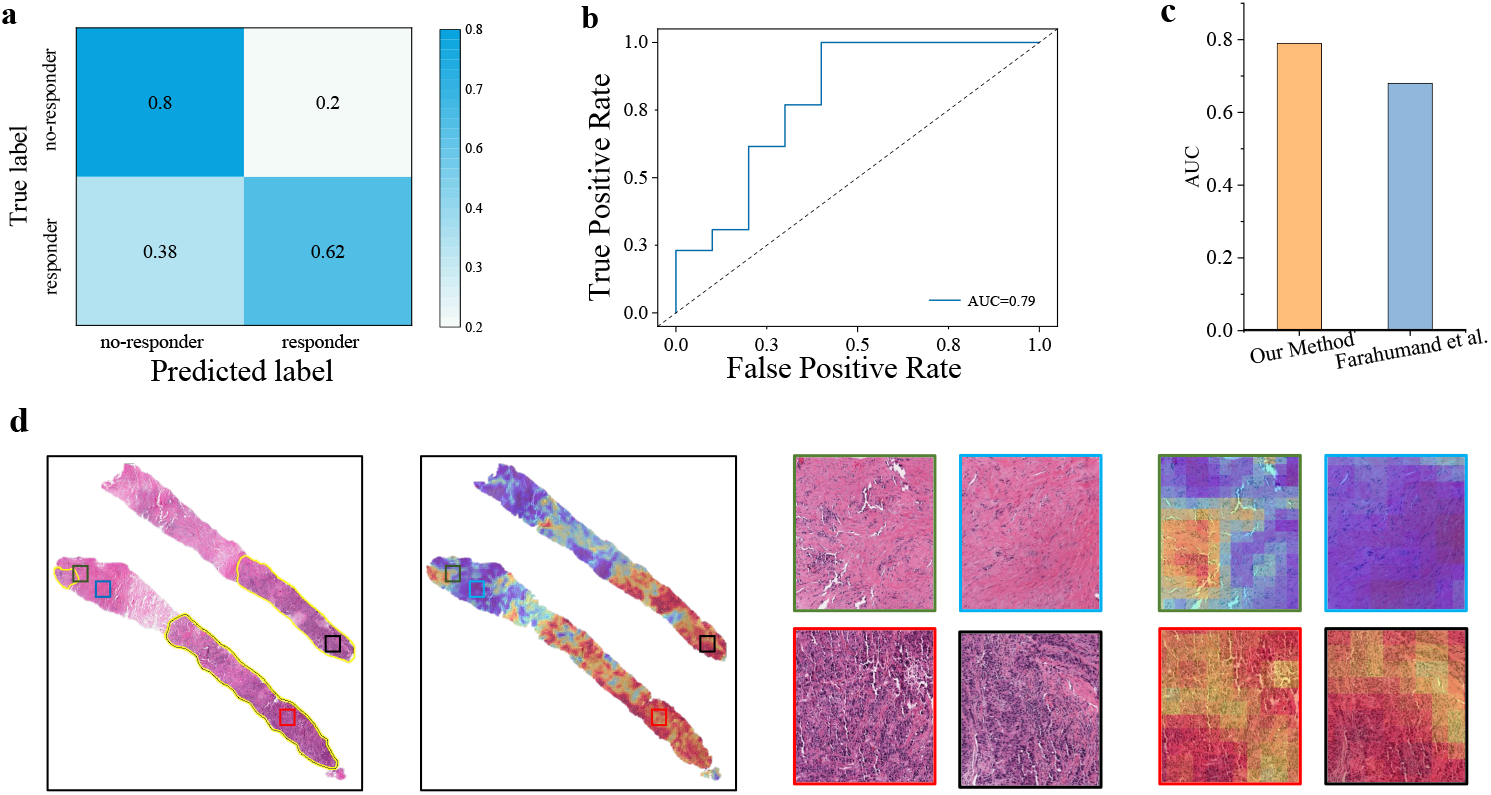
**(a-b)** showed the confusion matrix and ROC curve of the predicted drug response on the Yale trastuzumab response cohort. **(c)** shows the AUC values achieved by our method and Farahmand et al. [31]. **(d)** showed two pathology images with labeled tumor regions by a Yale pathologist and the heatmaps generated by our model (left), as well as some tiles randomly selected from the manually annotated regions (right). The tiles with black and red border represent tumor area, blue represents normal area, and green represents the junction area.

### 2.7 Spatial transcriptomics validates spatial localization of tumor biomarkers

Spatial transcriptomics measures the RNA abundance at a high spatial resolution enable us to evaluate our model capacity in spatial localization of tumor biomarkers. For this purpose, we used the RPPA-measured protein levels of two typical biomarkers, HER2 and TP53, to fine-tune our model. Next, we generated the heatmaps using the learned tile-level attention scores. Correspondingly, we visualized the spatial expression of HER2 and TP53 genes. As shown in Figure 7, these heatmaps showed that the highly-scored regions were notably consistent with the high expression region of the spatial transcription for both HER2 and TP53. This finding showed that our method effectively performed spatial deconvolution of tumor biomarkers based on conventional H&E staining sections.

**Fig. 7:**
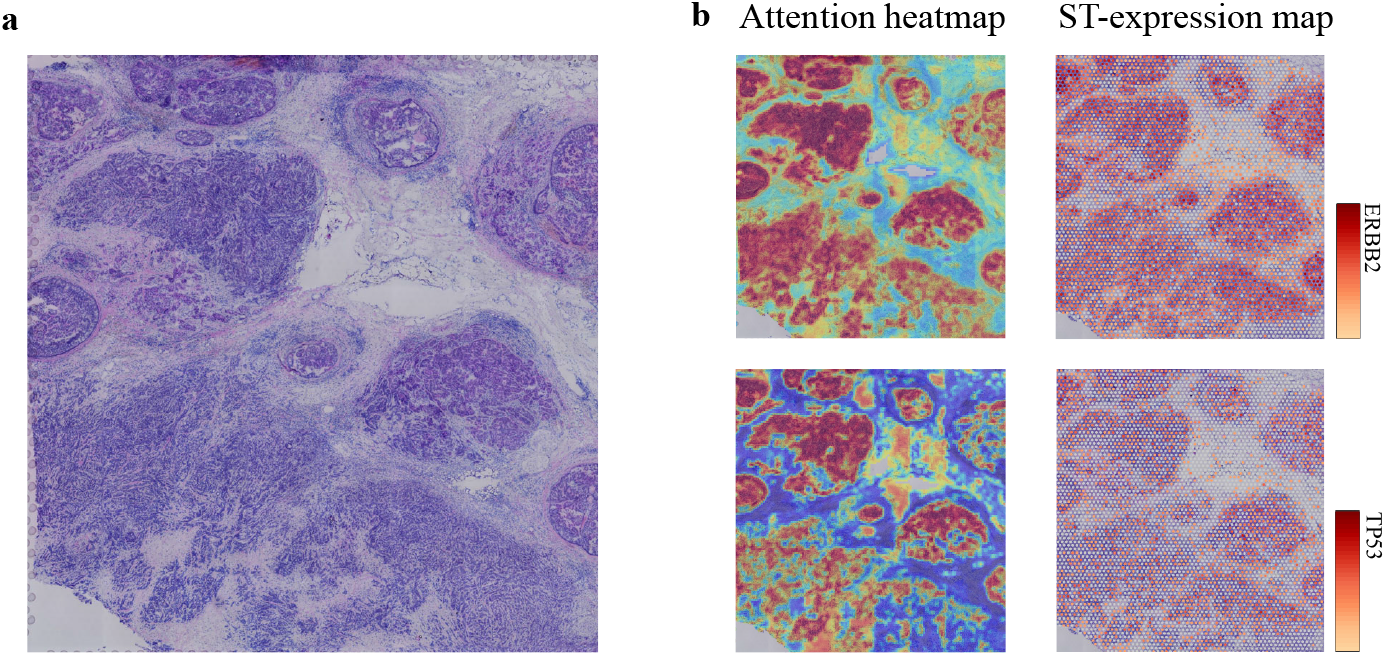
**(a)** Original pathological section used for spatial transcriptome sequencing. **(b)** Heatmaps showing the ERBB2 and TP53 spatially expressed landscapes generated by our model and spatial transcriptomic data.

## 3 Materials and methods

### 3.1 Whole slide images

All digital slides of fresh frozen tissue stained with hematoxylin and eosin (H&E) were obtained from TCGA via the Genomic Data Commons Data Portal. From the TCGA-BRCA project, we collected 1,979 WSIs of 1,094 breast cancer patients, comprising 1,580 tumor slides and 399 normal slides. The pathology diagnosis provided by TCGA database were used as ground truth labels for classification task.

An independent cohort came from Yale trastuzumab response cohort was used to evaluate model ability in predicting drug response. The Yale cohort contained 75 FFPE WSIs of 75 cases of breast cancer (34 responder and 41 non-responders). A case was designated as responder if the pathological examination of surgical resection specimens did not report residual invasive, lympho-vascular invasion or metastatic carcinoma, and otherwise non-responders. Note that only the slides with a magnification greater than 20* were included in this study.

### 3.2 RPPA dataset

All RPPA data were downloaded from The Cancer Proteome Atlas (TCPA)[32]. The database collects the RPPA-assayed protein levels of samples mainly came from TCGA (The Cancer Genome Atlas) tumor tissue sample sets. The RPPA data contains more than 200 proteins covering most cancer signaling pathways such as PI3K, MAPK, mTOR, TGF-b and WNT pathways. In total, there are 893 RPPA samples came from TCGA-BRCA cohort. After removal of sample without matched WSIs, we obtained 860 matched samples that are used to train protein level prediction model.

### 3.3 Spatial transcriptomic data

We acquired the spatial transcriptomic data of a breast cancer specimen using the Visium Spatial Gene Expression protocol from website. To explore the spatial map of gene expression, we utilized the 10x Loupe Browser, a desktop application that offers interactive visualization capabilities for various 10x Genomics solutions. This enabled us to visualize the spatial landscape of specific biomarker genes experssion upon the pathology image.

### 3.4 Preprocessing of whole slide images

Due to the ultra-high resolution of pathological images, which can reach up to gigapixels, they are not immediately suitable for input into a deep learning model. As a result, we divided each whole slide image into small squares known as tiles or patches. The Python package OpenSlide was utilized to read a slide into memory, and the Otsu algorithm was then employed to differentiate between tissue and background regions. Following segmentation, the tissue area was divided into 256*256 px tiles.

### 3.5 Contrastive learning for feature extraction

Given a large scale of unlabeled tiles, we leveraged the contrastive learningbased pretraining to learn an encoder to produce latent representations for the tiles. In this study, we used MoCo v2 for the pretraining. We also tested another contrastive learning method, SimCLR, and found that MoCo v2 achieved better performance in downstream tasks.

Formally, contrastive learning learns to extract feature by minimizing the distance between the representations of positive pairs (e.g. image and its augmentation) and maximizing the distance between representations of negative pairs (e.g. different images). Suppose an tile *x* has its augmentation *x*^*′*^. We use a CNN network as the backbone encoder *f* to obtain their embedding *h* and *h*^*′*^, where *h* = *f* (*x*). Subsequently, *h* goes through a projection head *g* composed of two fully connected layers to produce vector *z* and *z*^*′*^, where *z* = *g*(*h*).

The contrastive loss in a minibatch was defined as follows:

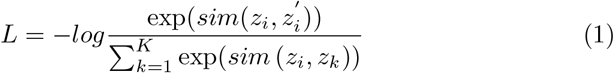

in which *sim*() is the similarity function, and *K* is the number of negative samples cached in memory bank. We adopted a simple similarity defined as below:

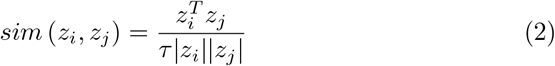

where *τ* is an adjustable temperature parameter. Once the pretraining was finished, we discarded the projection head and used the trained encoder to obtain the feature of tiles.

We used ResNet50 [33] as the backbone network, and loaded the pretrained weights on ImageNet. The SGD optimizer was used, learning rate of the backbone network was set to 0.03, the weight decay rate was 0.0001, and the momentum was 0.9.

### 3.6 Feature aggregation

For downstream tasks, we need to aggregate individual tile features into slide-level features. Instead of traditional max-pooling or average-pooling, we employ gated-attention pooling[34]. Suppose a slide has *N* tiles S= {*p*_1_, *p*_2_, …*p*_*N*_}, we got the feature *h*_*i*_ using the pretrained encoder for tile *p*_*i*_, the gated-attention pooling is essentially the instance-level weighted average pooling:

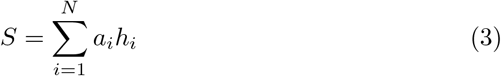

and

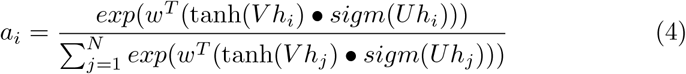

where U and V are trainable parameter matrices, • denotes element-wise multiplication, *sigm*() is a sigmoid non-linear activation function. The introduction of learnable parameters that are updated iteratively during the training process makes the model highly flexible and adaptable to various downstream tasks. Furthermore, after training, the attention weights reflect the significance of tile-level features for the downstream task, rendering the model interpretable.

### 3.7 Tumor diagnosis

We formulated the tumor diagnosis as the binary classification task. The ground truth labels (tumor or normal) came from TCGA-BRCA cohort. We employed a fully-connected layer plus a softmax layer as the prediction model that took the slide feature as input. The cross-entropy was used as the loss function for the classification task.

### 3.8 Protein level prediction

The protein level estimation for 223 tumor biomarkers was formulated as multi-task regression task. Taking as input the slide features, we adopted a multi-task learning model with a fully-connected layer and a output layer. The output layer had 223 nodes and each of them correspond to the level of a protein. In our practice, we have also tested multiple hidden layers architecture, and found that a single hidden layer could achieve superior performance in the regression task. The mean squared error (MSE) was used as loss function:

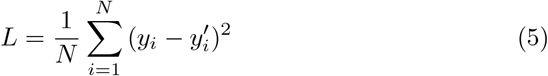

Where *y*_*i*_ is the measured level of protein *i* by RPPA assays, 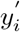 is the predicted level, and *N* is the total number of proteins.

### 3.9 Drug response prediction

The Yale trastuzumab response cohort was used to evaluate the model capacity to predict drug response. As a case was designated as responder or non-responders, we formualted the prediction of drug response from pathological features as binary classification problems, and the cross-entropy loss function was used.

### 3.10 Prognosis model

We used slide-level features to predict prognostic risk for each breast cancer patient. Given the slide feature *x*, time period *T*, and an event indicator *E*, we used a Cox proportional hazard regression model based on deep learning to estimate prognostic risk. The hazard function is defined as:

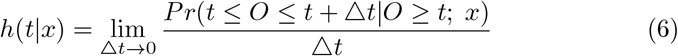

s It estimates the instantaneous death rate of individual *x* at time *t*. The Cox proportional hazard regression models it as:

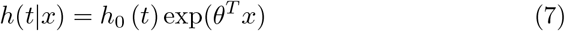

where *h*_0_ represents the baseline risk function at time *t*, which is estimated by Breslow estimation method. Our prognostic model was constructed and trained using the Pycox Python package. During training, the follow-up status and survival time are used to compute censored data and risk set. To estimate the parameters *θ*, we minimized the negative log partial likelihood:

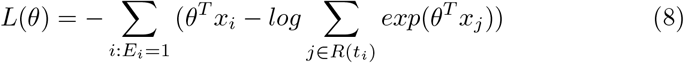

*R*(*t*_*i*_) is the risk set of time *t*_*i*_, which represents the set of patients who may be at risk at time *t*, and *E*_*i*_ = 1 indicates the death event.

## 4 Discussion and Conclusion

Proteomics plays a crucial role in cancer research and translational medicine by identifying key biomarkers for treatment and prognosis. Variations in protein expression levels are indicative of alterations in gene expression, which correspond to changes in cell and tissue phenotypes. By examining these variations, researchers can gain insights into the underlying molecular mechanisms of cancer and develop more effective diagnostic and treatment strategies. Proteomics also has the potential to improve the accuracy of personalized medicine by providing more comprehensive information about an individual unique molecular profile. With the rapid advancement of computational pathology, it has been found that histopathological features are predictive of gene mutations and microsatellite instability. However, few studies have focused on inferring protein levels from pathology images.

While comprehensive molecular tests, such as immunohistochemistry, are difficult to cover large-scale biomarkers, tissue sections stained with hematoxylin and eosin are ubiquitously available. Therefore, we set about to predict the protein levels of tumor biomarkers via computational pathology. We hypothesized that these routine tissue sections contain information about established and candidate biomarkers, so that molecular biomarkers could be inferred directly from digitized whole-slide images (WSIs). The rationale for this hypothesis is that protein level changes in cells cause changes of cellular function, which influence cell morphology. The underlying molecular profiles eventually dominate the histological characteristics, resulting in higher-order genotype–phenotype correlations. In fact, our experiments have demonstrated that pathological features faithfully reflected the protein levels in the specimen. There is increasing evidence that solid tumors exhibit significant tumor heterogeneity, namely, each tumor cell has distinct molecular genetic and phenotypic characteristics. This heterogeneity results in differences in growth rate, invasiveness, and drug sensitivity among tumor cells. Targeted drugs can only kill cells expressing specific target proteins, while a small number of drug-resistance tumor cells survive and proliferate, leading to tumor recurrence and progression. Although single-cell sequencing provides high-resolution data to reveal tumor heterogeneity, its high cost and long turnaround time hinder its widespread adoption in routine clinical examinations. In this work, whole slide images were divided into a large number of small tiles, each assigned a weight reflecting the importance of molecular expression levels within different regions of the tumor. This actually provides a fast and inexpensive alternative method for exploring tumor heterogeneity. Although current method based on pathological images has not reached clinically applicable standards, we believe that with the accumulation of data, especially the development of spatial transcriptomics, new computational pathology-based methodology will emerge to provide new insights into tumor heterogeneity from a spatial perspective.

In summary, we proposed a weakly supervised contrastive learning framework to infer protein levels of tumor biomarkers from breast cancer whole-slide images (WSIs). By pre-training the model on large-scale unlabeled breast cancer WSIs, the computational pathological features showed remarkable performance in various downstream tasks. The method performed well in tumor diagnosis and achieved high performance in predicting clinically relevant protein levels. The model interpretability is demonstrated through spatial visualization of WSIs colored by attention scores. Especially, our method also achieved notably accuracy in predicting the response of breast cancer patients to the drug trastuzumab.

## Declarations

### Data Availability

The whole slide images and corresponding labels of the TCGA-BRCA cohort from the TCGA database are available at https://portal.gdc.cancer.gov/. All RPPA data from The Cancer Proteome Atlas (TCPA) database is available at https://www.tcpaportal.org/tcpa/. The whole slide images and drug response data from Yale trastuzumab response cohort are available at TCIA database https://wiki.cancerimagingarchive.net/. The spatial transcriptomic data of a breast cancer specimen from 10x genomics are available at https://www.10xgenomics.com/. All other data supporting the findings of this study are available from the corresponding author upon reasonable request. Source data are provided with this paper.

### Code availability

All code was implemented using PyTorch as the primary deep-learning library. The complete pipeline for processing WSIs as well as training and evaluating our model are available at https://github.com/hliulab/wsi2rppa.

### Authors’ contributions

H.L. and X.X. conceived the study and designed the experiments. X.X. performed the experimental analysis. H.L. and X.X. prepared the manuscript. B.W. help to annotate the slides and supervised the research.

